# Brain network dynamics correlate with personality traits

**DOI:** 10.1101/702266

**Authors:** Aya Kabbara, Veronique Paban, Arnaud Weill, Julien Modolo, Mahmoud Hassan

## Abstract

**Introduction:** Identifying the neural substrates underlying the personality traits is a topic of great interest. On the other hand, it is now established that the brain is a dynamic networked system which can be studied using functional connectivity techniques. However, much of the current understanding of personality-related differences in functional connectivity has been obtained through the stationary analysis, which does not capture the complex dynamical properties of brain networks.

**Objective:** In this study, we aimed to evaluate the feasibility of using dynamic network measures to predict personality traits.

**Method:** Using the EEG/MEG source connectivity method combined with a sliding window approach, dynamic functional brain networks were reconstructed from two datasets: 1) Resting state EEG data acquired from 56 subjects. 2) Resting state MEG data provided from the Human Connectome Project. Then, several dynamic functional connectivity metrics were evaluated.

**Results:** Similar observations were obtained by the two modalities (EEG and MEG) according to the neuroticism, which showed a negative correlation with the dynamic variability of resting state brain networks. In particular, a significant relationship between this personality trait and the dynamic variability of the temporal lobe regions was observed. Results also revealed that extraversion and openness are positively correlated with the dynamics of the brain networks.

**Conclusion:** These findings highlight the importance of tracking the dynamics of functional brain networks to improve our understanding about the neural substrates of personality.

## Introduction

Personality refers to a characteristic way of thinking, behaving and feeling, that distinguishes one person from another (Back, Schmukle, & Egloff, 2009; Furr, 2009; Hong, Paunonen, & Slade, 2008; Jaccard, 1974). Since personality traits are thought to be stable and broadly predictable (Canli & Amin, 2002; Deyoung, 2006), it is unsurprising that personality is linked to reliable markers of brain function (Yarkoni, 2014). In this context, the interest in the neural substrates underpinning personality has substantially increased in recent years. One of the most widely used and accepted taxonomies of personality traits is the factor five model (FFM), or big-five model, which covers different aspects of behavioral and emotional characteristics (McCrae & John, 1992). It represents five main factors: conscientiousness, openness to experience, neuroticism, agreeableness and extraversion.

On the other side, emerging evidence shows that most cognitive states and behavioral functions depend on the activity of numerous brain regions operating as a large-scale network (Bressler, 1995; Edelman, 1993; Fuster, 2010; Goldman-Rakic, 1988; Greicius, Krasnow, Reiss, & Menon, 2003; Mesulam, 1990; O Sporns, Chialvo, Kaiser, & Hilgetag, 2004). This dynamical behavior is even present in the pattern of intrinsic or spontaneous brain activity (i.e., when the person is at rest) (Allen et al., 2014; Baker et al., 2014; F. de Pasquale, Penna, Sporns, Romani, & Corbetta, 2015; Francesco de Pasquale et al., 2012; Kabbara, Falou, Khalil, Wendling, & Hassan, 2017a; O’Neill et al., 2017). In particular, the dynamics of brain connectivity patterns can be studied at the millisecond time scale, for example using electro-encephalography (EEG) and magneto-encephalography (MEG).

However, while multiple studies have been conducted to relate the FFM traits to functional patterns of brain networks (Beaty et al., 2016a; Li et al., 2017; Mulders, Llera, Tendolkar, van Eijndhoven, & Beckmann, 2018; Tian, Wang, Xu, Li, & Ma, 2018; Tomecek & Androvicová, 2017; Toschi, Riccelli, Indovina, Terracciano, & Passamonti, 2018), we argue that the assessment of such relationships has been limited, in large part, due to an ignorance of networks variation throughout the measurement period. In the present study, we hypothesized that investigating the dynamic properties of the brain network reconfiguration over time will reveal new insights about the neural substrate of personality. Our hypothesis was supported by many recent studies that demonstrate the importance of examining the temporal variations of brain networks in personality traits such as intelligence, creativity, executive function and resilience (Kenett, Betzel, & Beaty, 2020; Tompson, Falk, Vettel, & Bassett, 2018; Paban, Modolo et al., 2020).

Here, we tested our hypothesis on two datasets: 1) Resting-state EEG data acquired from 56 subjects, and 2) Resting-state MEG data provided from the publicly available Human Connectome Project (HCP) MEG2 release including 61 subjects. Dynamic brain networks were reconstructed using the EEG/MEG source connectivity approach (Hassan & Wendling, 2018) combined with a sliding window approach as in (Kabbara et al., 2017a; O’Neill et al., 2017; Rizkallah et al., 2018). Then, based on graph theoretical approaches, several dynamic features were estimated. Correlations between individual FFM traits and network dynamics were assessed. Our findings reveal robust relationships between dynamic network measures and four of the big five personality traits (openness, conscientiousness, extraversion and neuroticism).

## Materials and methods

The full pipeline of the current study is summarized in Figure 1.

**Figure 1.**
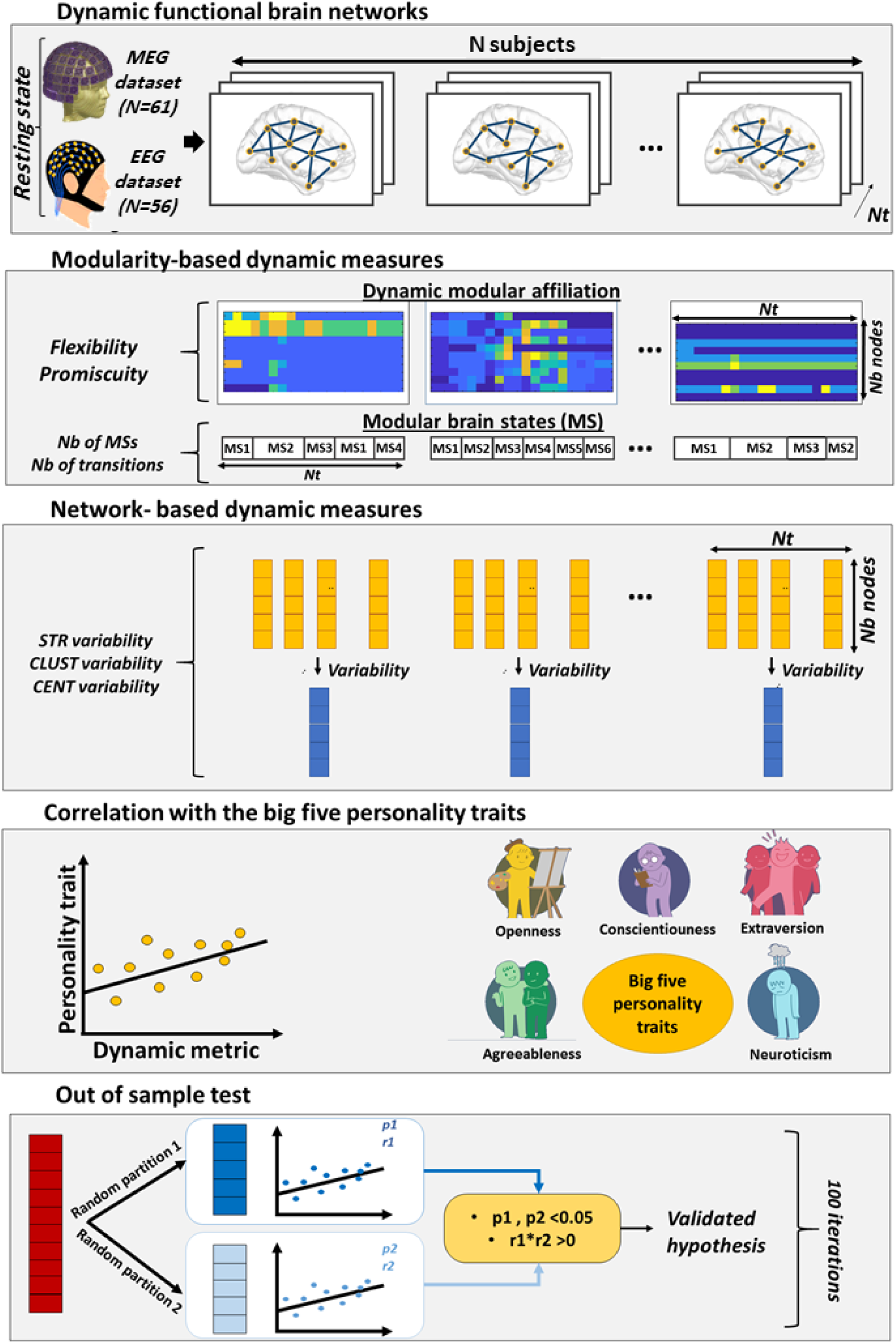
Full study pipeline. First, dynamic brain networks were reconstructed from resting state EEG data of 56 participants and MEG data of 61 participants. Then, for each subject, dynamic features were extracted (modularity-based features and graph-based features). Correlations between FFM personality traits (agreeableness, extraversion, neuroticism, openness, conscientiousness) and the dynamic features were then evaluated. Finally, statistical tests were assessed using a randomized out of sample test. STR = strength, CLUST=clustering coefficient, CENT=betweenness centrality.

### Dataset 1: EEG dataset

#### Participants

A total of 56 healthy subjects were recruited (29 women). The mean age was 34.7 years old (*SD* = 9.1 years, range = 18–55). Education ranged from 10 years of schooling to a PhD degree. None of the volunteers reported taking any medication or drugs, nor suffered from any past or present neurological or psychiatric disease. The study was approved by the “Comité de Protection des Personnes Sud Méditerranée” (agreement n° 10–41).

#### EEG Acquisition and Preprocessing

Each EEG session consisted in a 10-min resting period with the participant’s eyes closed (Paban, Deshayes, Ferrer, Weill, & Alescio-Lautier, 2018). Participants were seated in a dimly lit room, were instructed to close their eyes, and then to simply relax until they were informed that they could open their eyes. Participants were informed that the resting period would last approximately 10 min. The eyes-closed resting EEG recordings protocol was chosen to minimize movement and sensory input effects on electrical brain activity. EEG data were collected using a 64-channel Biosemi ActiveTwo system (Biosemi Instruments, Amsterdam, The Netherlands) positioned according to the standard 10–20 system montage, one electrocardiogram, and two bilateral electro-oculogram electrodes (EOG) for horizontal movements. Nasion-inion and preauricular anatomical measurements were made to locate each individual’s vertex site. Electrode impedances were kept below 20 kOhm. EEG signals are frequently contaminated by several sources of artifacts, which were addressed using the same preprocessing steps as described in several previous studies dealing with EEG resting-state data (Kabbara et al., 2017; Kabbara et al., 2018; Rizkallah et al., 2018). Briefly, bad channels (signals that are either completely flat or contaminated by movement artifacts) were identified by visual inspection, complemented by the power spectral density. These bad channels were then recovered using an interpolation procedure implemented in Brainstorm (Tadel, Baillet, Mosher, Pantazis, & Leahy, 2011) by using neighboring electrodes within a 5-cm radius. Epochs with voltage fluctuations between +80 μV and −80 μV were kept. Five artifact-free epochs of 40-s length were selected for each participant. This epoch length was used in a previous study, and was considered as a good compromise between the needed temporal resolution and the results reproducibility (Kabbara et al., 2017a).

#### Dynamic brain networks construction

Dynamic brain networks were reconstructed using the “EEG source connectivity” method (M Hassan & Wendling, 2018), combined with a sliding window approach as detailed in (Kabbara et al., 2017; Kabbara et al., 2018; Rizkallah et al., 2018). “EEG source connectivity” involves two main steps: i) solving the inverse problem in order to estimate the cortical sources and reconstruct their temporal dynamics, and ii) measuring the functional connectivity between the reconstructed time-series.

Briefly, the steps performed were the following:

1. EEGs and MRI template (ICBM152) were coregistered through the identification of anatomical landmarks by using Brainstorm (Tadel et al., 2011).
2. A realistic head model was built using the OpenMEEG (Gramfort, Papadopoulo, Olivi, & Clerc, 2010) software.
3. A Desikan-Killiany atlas-based segmentation approach was used to parcellate the cortical surface into 68 regions (Desikan et al., 2006).
4. The weighted minimum norm estimate (wMNE) algorithm was used to estimate the regional time series (Hamalainen & Ilmoniemi, 1994).
5. The reconstructed regional time series were filtered in different frequency bands (delta: 1–4 Hz; theta: 4–8 Hz; alpha: 8–13 Hz; beta: 13–30 Hz and gamma: 30-45 Hz)
6. To compute the functional connectivity between the reconstructed regional time-series, we used the phase locking value (PLV) metric (Lachaux et al., 2000) defined by the following equation:

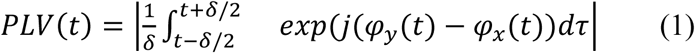

where *φ*_*y*_(*t*) and *φ*_*x*_(*t*) are the unwrapped phases of the signals *x* and *y* at time *t*. The Hilbert transform was used to comput the instantaneous phase of each signal. *δ* denotes the size of the window in which PLV is calculated. Dynamic functional connectivity matrices were computed for each epoch using a sliding window technique (Kabbara, Falou, Khalil, Wendling, & Hassan, 2017b). It consists in moving a time window of certain duration *δ* along the time dimension of the epoch, and then PLV is calculated within each window. As recommended by (Lachaux et al., 2000), the number of cycles should be sufficient to estimate PLV in a compromise between a good temporal resolution and a good accuracy. The smallest number of cycles recommended equals to 6. In each frequency band, we chose the smallest window length that is equal to 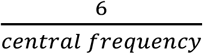. Thus, in delta band, as the central frequency (Cf) equals to 2.5 Hz, *δ* equals 2.4 s. Likewise, *δ* = 1s in the delta band (Cf=6 Hz), 571 ms in the alpha band (Cf=10.5 Hz),279 ms (Cf=21.5 Hz) in the beta band, 160 ms (Cf-37.5 Hz) in the gamma band. Functional connectivity matrices were represented as graphs (i.e networks) composed of nodes, represented by the 68 ROIs, and edges corresponding to the functional connectivity values computed over the 68 regions, pair-wise.
7. To ensure equal network density for all the dynamic networks computed across time, a proportional (density-based) threshold was applied in a way to keep the top 10% of connectivity values in each network.

### Dataset 2: MEG dataset *(HCP)*

#### Participants

As part of the HCP MEG2 release (Larson-Prior et al., 2013; Van Essen et al., 2012), resting-state MEG recordings were collected from 61 healthy subjects (38 women). The release included 67 subjects, but six subjects were omitted from the analysis as their recordings failed to pass the quality control checks (including tests for excessive SQUID jumps, sensible power spectra, correlations between sensors, and availability of sufficient good quality recording channels). All subjects are young (22–35 years of age) and healthy.

#### MEG recordings and pre-processing

The acquisition was performed using a whole-head Magnes 3600 scanner (4D Neuroimaging, San Diego, CA, USA). Resting state measurements were taken in three consecutive sessions of 6 min each. Data were provided pre-processed, after passing through a pipeline that removed artefactual segments, identified faulty recording channels, and regressed out artefacts which appear as independent components in an ICA decomposition with clear artefactual temporal signatures (such as eye blinks or cardiac interference).

#### Dynamic brain networks construction

Here, we adopted the same pipeline used by the previous studies dealing with the same dataset (Colclough et al., 2016). Thus, to solve the inverse problem, we have applied a linearly constrained minimum variance beamformer (Van Veen, Van Drongelen, Yuchtman, & Suzuki, 1997). Pre-computed single-shell source models are provided by the HCP and the data covariance were computed separately in the 1–30 Hz and 30–48 Hz bands as in (Colclough et al., 2016). Data were beamformed onto a 6 mm grid using normalized lead fields. Then, source estimates were normalized by the power of the projected sensor noise. Source space data were filtered in delta: 1–4 Hz; theta: 4–8 Hz; alpha: 8–13 Hz; beta: 13–30 Hz and gamma: 30-45 Hz (as in EEG dataset). After obtaining the regional time series on the basis of the Desikan-Killiany atlas, a symmetric orthogonalization procedure (Colclough, Brookes, Smith, & Woolrich, 2015) was performed for signal leakage removal. To ultimately estimate the functional connectivity between regional time series, we used the amplitude envelope correlation measure (AEC) (M. J. Brookes, Woolrich, & Barnes, 2012). This method briefly consists of 1) computing the power envelopes as the magnitude of the signal, using the Hilbert transform, and 2) measuring the linear amplitude correlation between the logarithms of ROI power envelopes. Finally, a sliding window (length = 6 sec, step = 0.5 sec) was applied to construct the dynamic connectivity matrices. This sliding window has been previously used to reconstruct the dynamic networks derived from MEG data (O’Neill et al., 2016). Also, matrices were thresholded by keeping the strongest 10% connections of each network.

### Dynamic measures

While functional connectivity provides crucial information about how the different brain regions are connected, graph theory offers a framework to characterize the network topology and organization. In practice, many graph measures can be extracted from networks to characterize static and dynamic network properties. Here, we focused on measures quantifying the dynamic aspect of the brain networks/modules/regions and their reconfiguration over time.

#### Graph-based dynamic measures

Most previous studies attempt to average the graph measures derived from temporal windows (F. de Pasquale et al., 2015; Kabbara et al., 2017a). However, such strategy constrains the dynamic analysis. Distinctively, we aimed here at quantifying the dynamic variation of node’s characteristics inferred from graph measures (including strength, centrality and clustering). The graph measure’s variation (*GM*) of the node *i* across time windows is defined as:

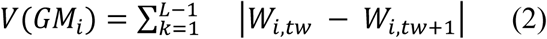

Where *GM* is the considered graph measure, *L* denotes the number of time windows and *tw* and *tw* + 1 refer to two consecutive time windows. *W*_*i,tw*_ is the value of the graph measure (strength, clustering or centrality) of the considered node *i* at the time window *tw*. A node with high V reflects that the node is dynamic in terms of the given *GM*. In this study we focused on three graph measures:

1. **Strength:** The node’s strength is defined as the sum of all edges weights connected to a node (Barrat, Barthélemy, Pastor-Satorras, & Vespignani, 2004). It indicates how influential the node is with respect to other nodes.
2. **Clustering coefficient:** The clustering coefficient of a node evaluates the density of connections formed by its neighbors (Watts & Strogatz, 1998). It is calculated by dividing the number of existing edges between the node’s neighbors to the number of possible edges. The clustering coefficient of a node is an indicator of its segregation within the network.
3. **Betweenness centrality:** The betweenness centrality calculates the number of shortest paths that pass through a specific node (Rubinov & Sporns, 2011). The importance of a node is proportional to the number of paths in which it participates.

An illustrative example of strength variability on a toy dynamic graph is presented in Figure 2.B.

**Figure 2.**
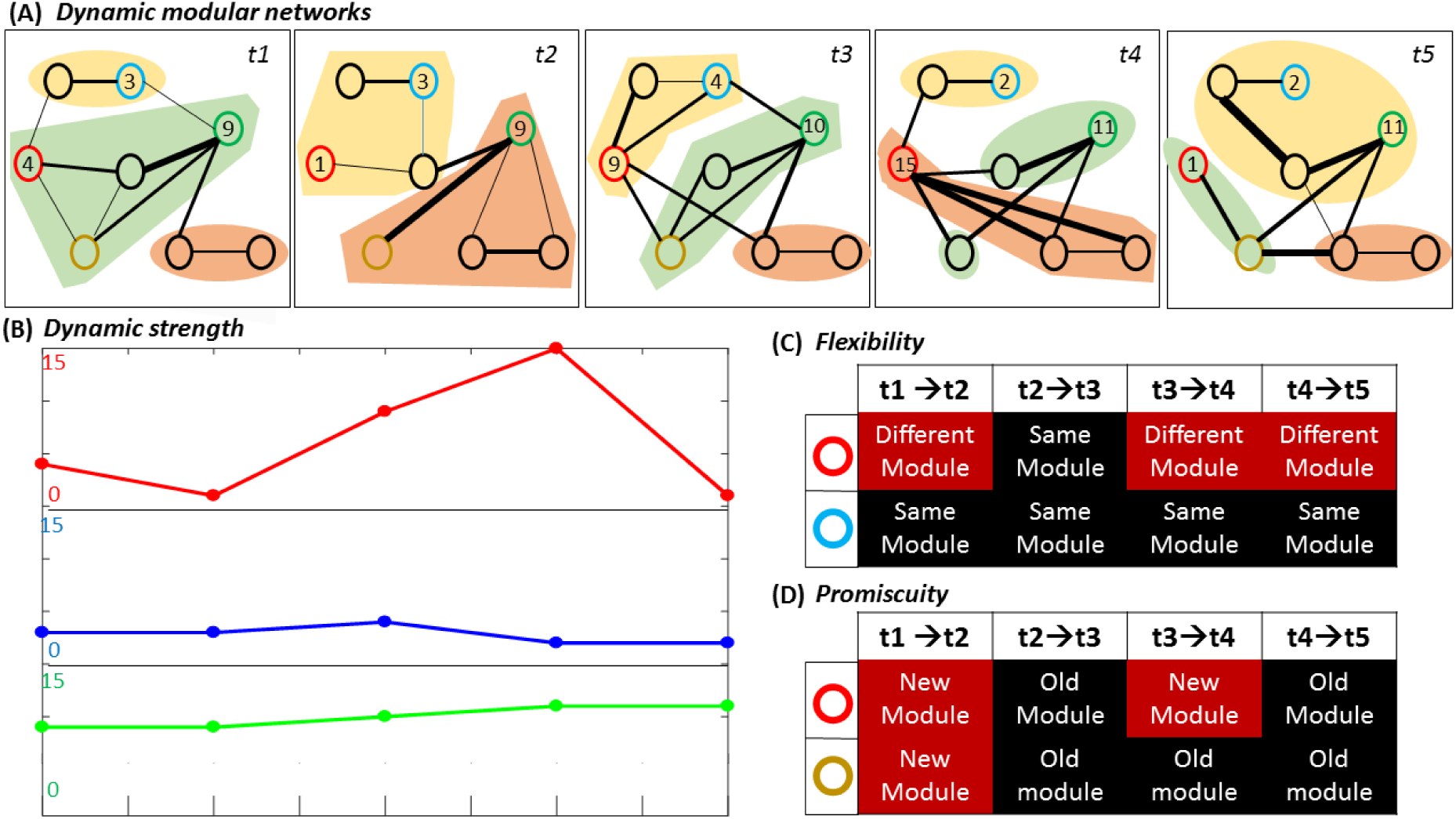
An illustrative example of the dynamic features extracted from a toy dynamic graph. A) The dynamic modular networks. B) The strength variation. C) Flexibility of red and blue nodes. D) Promiscuity of red and yellow nodes.

#### Modularity-based dynamic measures

Modularity describes the tendency of a network to be partitioned into modules or communities of high internal connectivity and low external connectivity (Sporns and Betzel, 2016). To explore how brain modular networks reshape over time, we detected the dynamic modular states that fluctuate over time using our recent proposed algorithm (Kabbara et al., 2019). Briefly, it attempts to extract the main modular structures (known as modular states) that fluctuate repetitively across time. Modular states reflect unique spatial modular organization, and are derived as follows:

- Decompose each temporal network into modules using the consensus modularity approach (Bassett et al., 2013; Kabbara et al., 2017a). This approach consists of generating an ensemble of partitions acquired from the Newman algorithm (Girvan & Newman, 2002) and Louvain algorithm (Blondel, Guillaume, Lambiotte, & Lefebvre, 2008) repeated for 200 runs. Then, an association matrix of N × N (where N is the number of nodes) is obtained by counting the number of times two nodes are assigned to the same module across all runs and algorithms. The association matrix is then compared to a null model association matrix generated from a permutation of the original partitions, and only the significant values are retained (Bassett et al., 2013). To ultimately obtain consensus communities, we re-clustered the association matrix using Louvain algorithm.
- Assess the similarity between the temporal modular structures using the z-score of Rand coefficient, bounded between 0 (no similar pair placements) and 1 (identical partitions) as proposed by (Traud, Kelsic, Mucha, & Porter, 2008). This yielded a T × T similarity matrix where T is the number of time windows.
- Cluster the similarity matrix into “categorical” modular states (MS) using the consensus modularity method. This step combines similar temporal modular structures into the same community. Hence, the association matrix of each “categorical” community is computed using the modular affiliations of its corresponding networks.

Once the modular states (MS) were computed, two metrics were extracted:

1. *The number of MSs*
2. *The number of transitions:* It measures the number of switching between MSs.

In addition, after obtaining the dynamic modular affiliations, two dynamic nodal measures were calculated:

1. *Flexibility:* It is defined as the number of times that a brain region changes its module across time, normalized by the total number of changes that are possible. We considered that a module was changed if more than 50% of its nodes have changed (Figure 2.C).
2. *Promiscuity:* It is defined as the number of modules a node participates during time (Figure 2.D)

### Statistical analysis

Dynamic measures were extracted at the level of each brain region (node-wise analysis), and at the level of the whole network. At the network-level, flexibility, promiscuity, strength variation, clustering variation and centrality variation were averaged over all brain regions. At the node-level, the values of each node were kept. In order to investigate the associations between the dynamic network measures and FFM personality traits, Pearson’s correlation analysis was assessed. To consider the multiple comparisons problem (between the five frequency bands, five personality traits and 68 ROIs), *p*-values were corrected using Bonferroni and FDR procedures (Bland & Altman, 1995). Bonferroni correction yields an adjusted threshold of *p* < 0.002 for the network-level. For node-level features, *p*-value were corrected across the five frequency bands, five personality traits and 68 regions, resulting in a Bonferroni-adjusted threshold of *p* < 2*E* − 5.

To avoid data dredging problem, we conducted randomized out-of-sample tests repeated 100 times. The out of sample test consists of randomly dividing data into two random subsets. If significant correlations were obtained from the two subsets for more than 95% of the iterations, the correlation is considered statistically significant on the whole distribution.

### Evaluating the FFM personality traits

The Five-Factor Model (FFM) represents five major personality traits: 1) *conscientiousness* which describes an organized and detailed-oriented nature, 2) *agreeableness* which is associated to kindness and cooperativeness, 3) *neuroticism* which indexes the tendency to have negative feelings, 4) *openness* is related to intellectual curiosity and imagination, 5) *extraversion* refers to the energy drawn from social interactions.

For the EEG dataset, personality traits were assessed with the French Big Five Inventory (BFI-Fr) (Plaisant, Courtois, Réveillère, Mendelsohn, & John, 2010). The BFI-Fr is composed by 45 items in which respondents decide whether they agree or disagree with each question, on a 1 (strongly disagree) to 5 (strongly agree) Likert scale. Responses are then summed to determine the scores for the five personality constructs.

According to the MEG dataset, the FFM personality traits were assessed via the NEO five-factors inventory (NEO-FFI) (Costa & McCrae, 1992; Terracciano, 2003). The NEO-FFI is composed by 60 items in which participants reported their level of agreement on a 5-points Likert scale, from strongly disagree to strongly agree.

## Results

In each dataset, the dynamic functional networks were reconstructed using a sliding window approach for each subject. Then, dynamic measures were extracted at the level of each brain region (node-wise analysis), and at the level of the whole network. At the network-level, flexibility, promiscuity, strength variation, clustering variation and centrality variation were averaged over all brain regions. At the node-level, the values of each node were kept.

### Dataset 1: EEG

The correlation between FFM personality traits and the network-level parameters are presented in Figure 3. Neuroticism showed a negative correlation with the number of transitions (*p*_*FDR corrected*_ < 0.05; *r* = −0.35) and the overall promiscuity (*p*_*FDR corrected*_ < 0.05; *r* = −0.44) in the beta band, as well as the flexibility in the theta band (*p*_*Bonferroni corrected*_ < 0.05; *r* = −0.37). Results also depict a negative correlation between conscientiousness and the overall clustering variation in the alpha band (*p*_*Bonferroni corrected*_ < 0.05; *r* = −0.43). No significant relationship was observed at the network-level between any of the dynamic measures with agreeableness, openness and extraversion.

**Figure 3.**
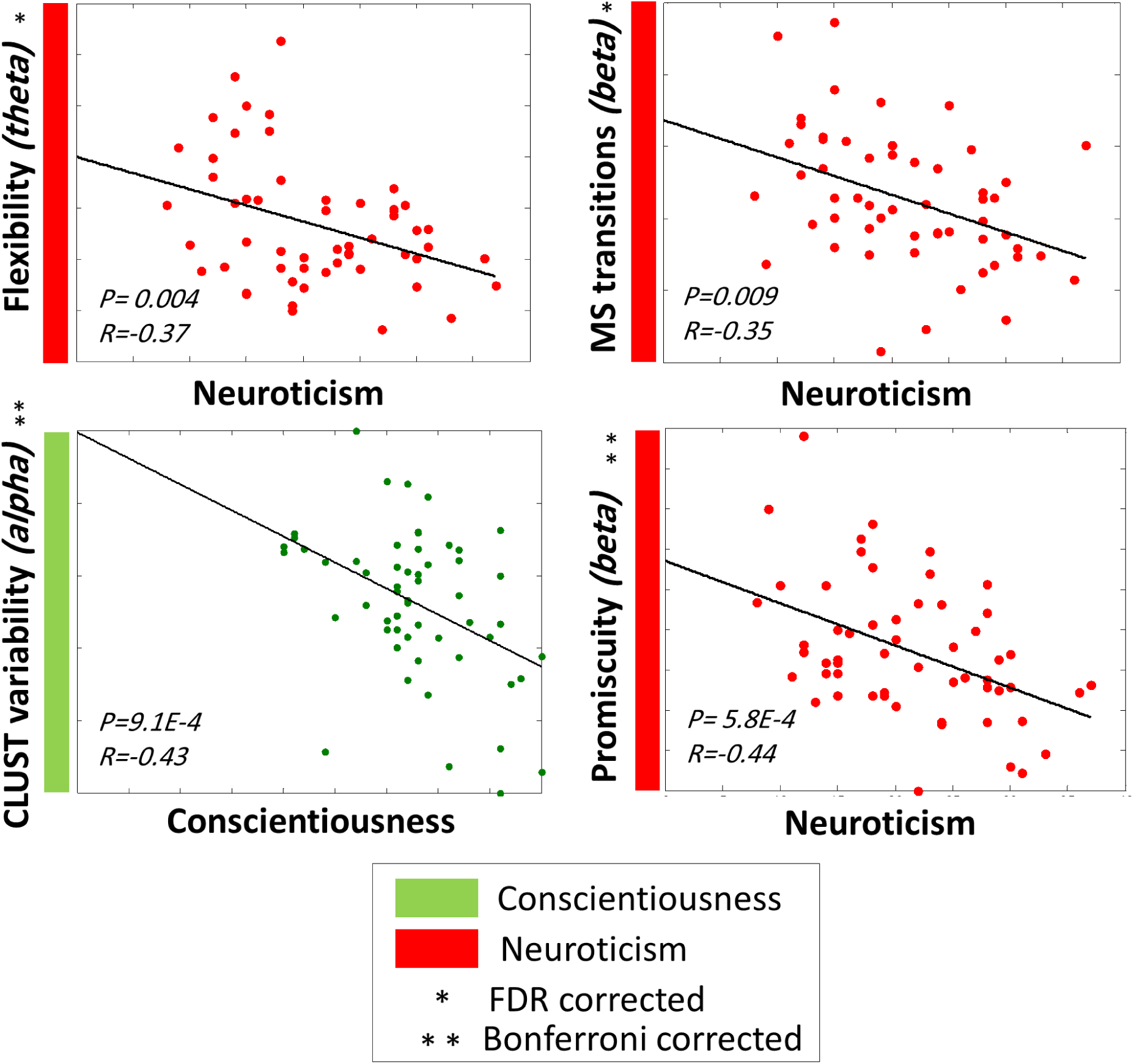
Significant correlations between the FFM traits and the dynamic graph measures computed on the network-level using EEG dataset.

Figure 4 illustrates the correlation between FFM traits and nodal characteristics in terms of dynamic features. Results show that higher extraversion was correlated with higher clustering variability of superior parietal lobule (SPL) in the theta band (*p*_*Bonferroni corrected*_ < 0.05; *r* = 0.54). In contrast, neuroticism was negatively correlated with strength variation of the left middle temporal gyrus (MTG) (*p*_*Bonferroni corrected*_ < 0.05; *r* = −0.51), left superior temporal gyrus (STG) (*p*_*Bonferroni corrected*_ < 0.05; *r* = −0.55) and transverse temporal gyrus (TT) (*p*_*FDR corrected*_ < 0.05; *r* = −0.5) in the theta band.

**Figure 4.**
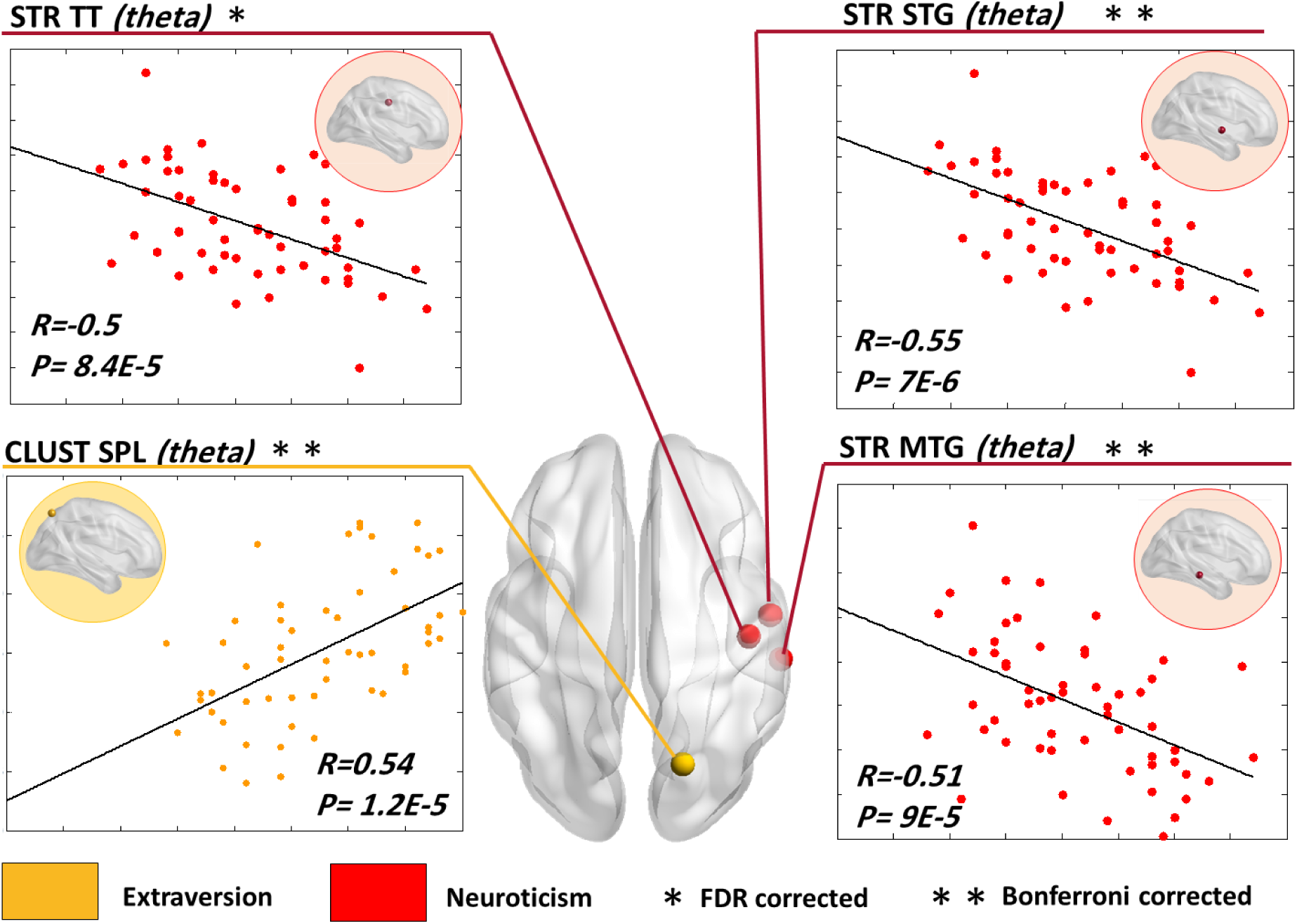
The cortical surface illustrating the brain regions for which the dynamic measures significantly correlated with FFM traits using EEG dataset. STR = strength, CLUST=clustering coefficient, CENT=betweenness centrality, FLEX: flexibility, FUS=Fusiform, PCC: posterior cingulate cortex SPL=superior parietal lobule, STG= superior temporal gyrus, MTG=middle temporal gyrus, TT= transverse temporal.

### Dataset 2: MEG

Figure 5 illustrates the correlation between FFM personality traits and network-level parameters for the MEG analysis. One can notice that neuroticism showed negative correlations with flexibility in the theta (*p*_*Bonferroni corrected*_ < 0.05; *r* = −0.36), alpha (*p*_*Bonferroni corrected*_ < 0.05; *r* = −0.46) and beta bands (*p*_*Bonferroni corrected*_ < 0.05; *r* = −0.39). Neuroticism was also negatively correlated with strength variability in delta band (*p*_*Bonferroni corrected*_ < 0.05; *r* = −0.39). In contrast, a positive significant correlation was depicted between extraversion and the clustering variability in the theta band (*p*_*FDR corrected*_ < 0.05; *r* = −0.35).

**Figure 5.**
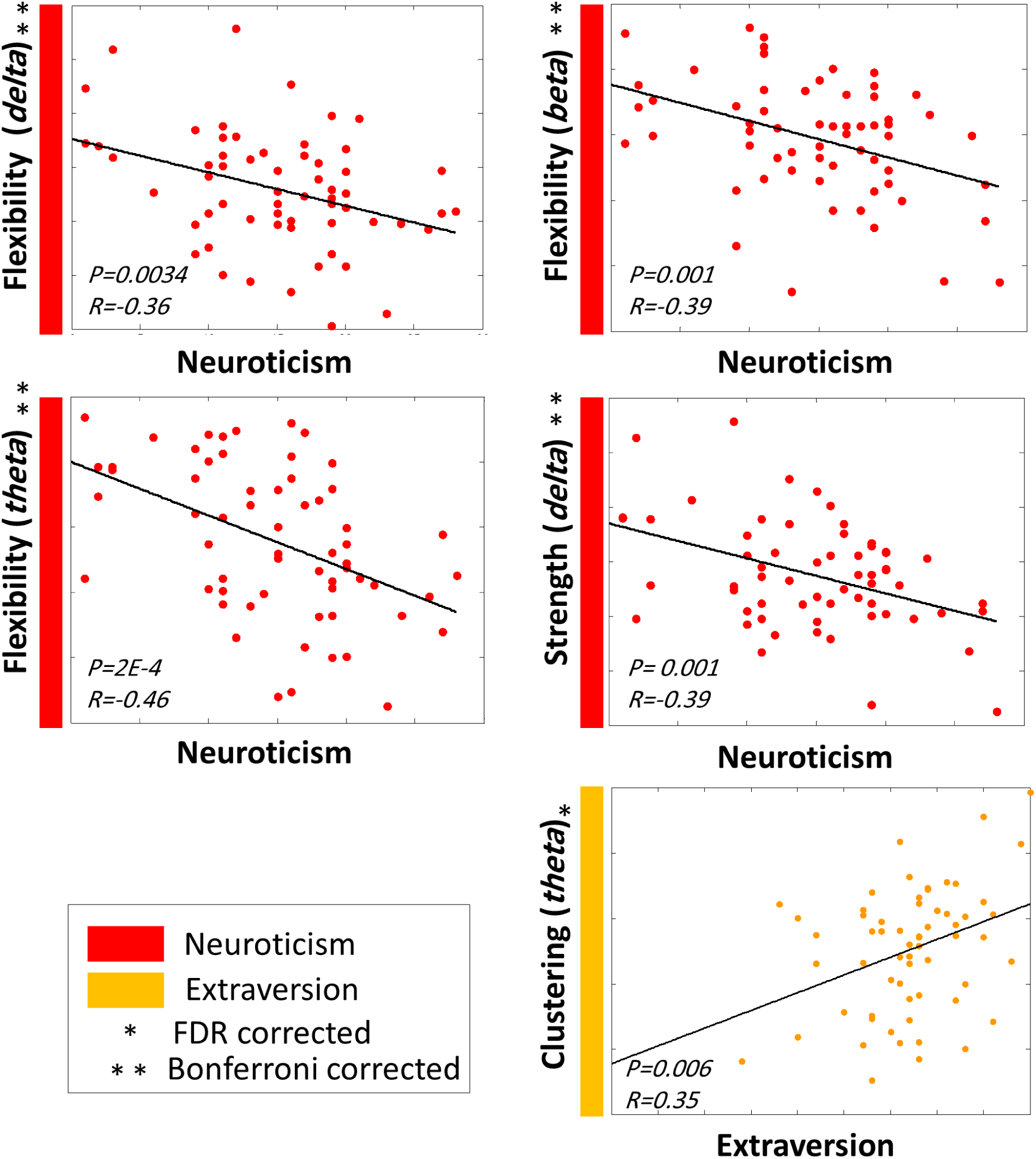
Significant correlations between the FFM traits and the dynamic graph measures computed on the network-level using MEG dataset.

Results in Figure 6 show that openness was positively correlated with the strength variability of the superior frontal gyrus (sFG) in the beta band (*p*_*FDR corrected*_ < 0.05; *r* = 0.48). However, negative correlations were observed between neuroticism and the strength variation of the left temporal pole (TP) in the alpha band (*p*_*Bonferroni corrected*_ < 0.05; *r* = −0.64), right supramarginal (SMAR) in both theta (*p*_*Bonferroni corrected*_ < 0.05; *r* = −0.54) and beta bands (*p*_*Bonferroni corrected*_ < 0.05; *r* = −0.54). In addition, neuroticism was negatively correlated with flexibility of the superior temporal gyrus (STG) in theta band (*p*_*Bonferroni corrected*_ < 0.05; *r* = −0.51).

**Figure 6.**
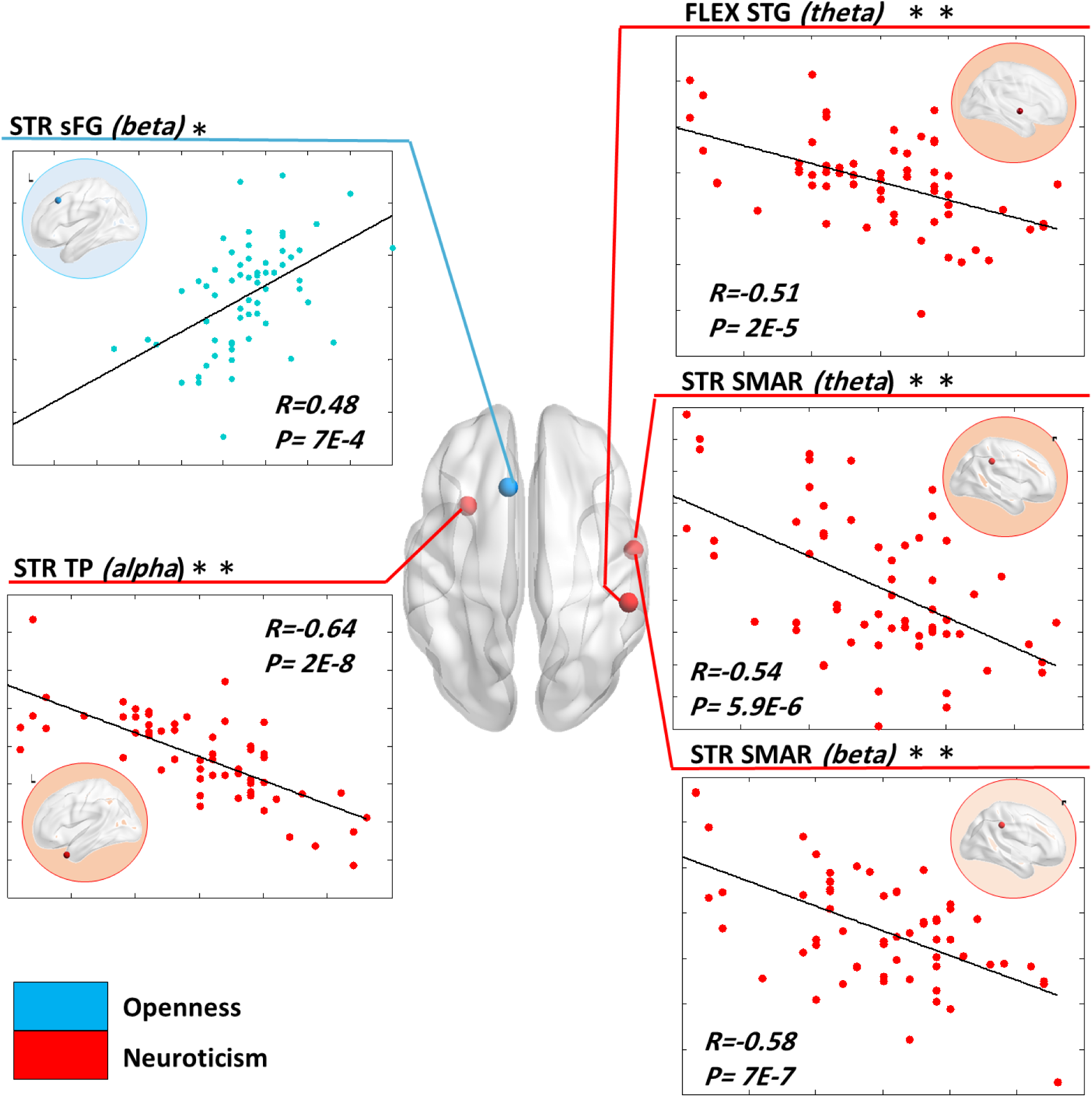
The cortical surface illustrating the brain regions for which the dynamic measures significantly correlated with FFM traits using MEG dataset. STR = strength, FLEX: flexibility, SMAR=Supramarginal, STG= superior temporal gyrus, TP= temporal pole.

### Randomized out of sample tests

For each feature, a distribution of 200 values (100 p-values for each random subset) was obtained as a result of the correlation between the FFM personality traits and the network feature. Figure 6.A shows a typical example of a node-level feature that successively passed the randomized tests. Specifically, the number of p-values lower than the Bonferroni adjusted value (***p* = 2*E* − 5**) reached 95% of the total number of iterations. In contrast, figure 6.B shows an example of a node-level feature that failed to pass the randomized tests with a proportion of 65% of significant correlations. We report in Table 1 and Table 2 the results of randomized tests for all features mentioned as significant for the two datasets.

## Discussion

The present study provides evidence that dynamic features (derived from graph measures) based on resting-state EEG data are significantly associated with FFM personality traits (derived from the BFI-Fr questionnaire).

The majority of studies in personality has mainly examined the interaction between neuropsychological traits and brain features in a static way. In particular, multiple previous studies focused on investigating how personality traits are linked to differences in morphological brain properties (DeYoung, 2010; Gray, Owens, Hyatt, & Miller, 2018; Liu et al., 2013; Omura, Constable, & Canli, 2005; Riccelli, Toschi, Nigro, Terracciano, & Passamonti, 2017). Another traditional way was to perform brain activation analysis to understand the neural basis of personality (Cooper, Tompson, O’Donnell, & Falk, 2015; Falk et al., 2015). However, these strategies ignore useful information about the way in which brain regions interact with each other (Sebastian Markett, Montag, & Reuter, 2018). Moving forward, multiple connectivity studies have been recently conducted to understand the neural substrates of human personality (Adelstein et al., 2011; Aghajani et al., 2013; Beaty et al., 2016b; Bey, Montag, Reuter, Weber, & Markett, 2015; Bey et al., 2015; Dubois, Galdi, Han, Paul, & Adolphs, 2018; Gao, 2013; Kyeong, Kim, Park, & Hwang, 2014; S. Markett et al., 2013; Sebastian Markett, Montag, Melchers, Weber, & Reuter, 2016; Tompson et al., 2018). Interestingly, graph theoretical assessment derived from networks was applied to link topological brain features to the Big Five personality traits (Beaty et al., 2016b; Bey et al., 2015; Gao, 2013; Toschi et al., 2018). As an example, (Toschi et al. 2018) shows that conscientiousness is linked to nodal properties (clustering coefficient, betweenness centrality and strength) of fronto-parietal and default mode network regions. Nevertheless, recent evidence revealed that dynamic analysis of functional data provides a more comprehensive understanding of neural implementation in personality (Tompson et al., 2018). The main originality of the current work is that it extends the traditional static view of brain networks to explore the time-varying characteristics associated to FFM traits. Particularly, we hypothesized that fast brain dynamics in EEG and MEG resting state networks are correlated with FFM personality. Our hypothesis is based on many recent studies suggesting that personality-related differences in functional connectivity are discernable during rest (Adelstein et al., 2011; Beaty et al., 2016; Bey, Montag, Reuter, Weber, & Markett, 2015; Gao, 2013; T. Li et al., 2017; Y. Li, Qin, Jiang, Zhang, & Yu, 2012; Markett et al., 2013; Mulders, Llera, Tendolkar, van Eijndhoven, & Beckmann, 2018; Sheu, Ryan, & Gianaros, 2011; Sheu et al., 2011). Such finding is advantageous since collecting brain data during rest is more feasible. Also, this hypothesis is supported by the evidence that resting-state brain dynamics fluctuates at sub-second timecale (less than 300 ms) (Baker et al., 2014; Damborská et al., 2019; Kabbara, Falou, Khalil, Wendling, & Hassan, 2017a).

At the level of the whole network, both EEG and MEG analyses showed common observations according to the neuroticism personality trait. This latter appeared to be the most sensitive to the analysis through dynamic approaches. Importantly, the EEG study showed negative correlations between neuroticism and centrality variation, number of transitions, promiscuity, and flexibility. Similarly, MEG study showed negative correlations between neuroticism, flexibility and strength variation. This suggests that the more individuals had a strong tendency to experience negative affection, such as anxiety, worry, fear, and depressive mood (Ormel et al., 2013), the less their brain showed dynamic characteristics in terms of modular organization over time. In other words, one may speculate that individuals with low dynamic measures of brain networks did not have enough capacity to get over their tendency to experience negative emotions and their psychological distress.

More particularly, at the node-level, the degree of neuroticism was associated with low dynamic variation of temporal regions using the two modalities (mainly STG, MTG and TT in EEG study; STG in MEG study). Importantly, the temporal lobe is known to be involved in processing sensory input related to visual memory, language comprehension, and emotion association (Kosslyn, 2007). In particular, the STG is involved in the interpretation of other individuals’ actions and intentions (Pelphrey & Morris, 2006). Others stated that STG plays an important role in emotional processing and effective responses to social cues, such as facial expressions and eye direction (Pelphrey & Carter, 2008; Singer, 2006). These findings are in agreement with a recent study showing that neurotic individuals present delayed detection of emotional and facial expressions (Sawada et al., 2016).

Using MEG dataset, extraversion was showed to be positively correlated with the clustering variation of the whole network. The similar dynamic behavior was also found using EEG dataset where a positive correlation was established between extraversion and the clustering variation of superior parietal lobule (SPL), which is involved in attention and visuomotor integration (Iacoboni & Zaidel, 2004). These findings highlight the complementary information that can be provided by the two modalities (F. de Pasquale, Corbetta, Betti, & Della Penna, 2018). In line with (Suslow et al., 2010) showing that extraverts displayed enhanced sensitivity and efficiency in sensory information processing compared with introverts, our data add to our neurobiological underpinning knowledge of extraversion highlighting the involvement of the SPL in such processes. Thus, SPL would play a central role promoting segregation within the network of extraverted individuals.

Besides these similar observations led by both MEG and EEG analyses, conscientiousness revealed a significant correlation with dynamic metrics only using EEG, while openness showed a significant correlation with the dynamic measures using MEG solely. This discrepancy can be due to the fact that MEG-EEG differences particularly arise when investigating the transient resting-state functional connectivity patterns (Coquelet et al., 2020). It may also be due to the difference in the sample analyzed by the two modalities, as well as the pre-processing, source reconstruction and connectivity methods used to reconstruct underlying networks. Moreover, several studies show that openness to experience and conscientiousness traits appear to differ across different samples (Hofstee, de Raad, & Goldberg, 1992; Johnson & Ostendorf, 1993). Still, the impact of these differences was less drastic on the neuroticism and the extraversion traits. Importantly, these two traits are universally accepted and appear in all major models of personality traits (Zelenski & Larsen, 1999). Thus, the most consistent and significant result obtained shows that the dynamic flexibility in functional networks could plausibly contribute to increased emotional reactivity, particularly linked to neuroticism and extraversion (Yarkoni, 2014).

Results show that among the five frequency bands studied, most changes were observed within slow oscillations (namely, delta, theta, and alpha bands). As suggested by (Knyazev, 2012), these oscillations might play a major role in integration across diverse cortical sites by synchronizing coherent activity and phase coupling across spatially distributed neural assemblies, so that it might not be surprising that network properties related to personality traits were affected only within slower frequency bands.

Overall, the present study adds to our recent paper (Paban et al. 2019) in providing new evidence that the dynamic reconfiguration of brain networks is of particular importance in shaping behavior.

### Limitations

In this study, we have assessed the personality traits using FFM. One common limitation of FFM is that it does not provide an adequate coverage of all personality domains (McAdams, 1992). As an example, it lacks the description of religiosity, honesty, sense of humor and many other domains. However, there is no consensus about the exact number of broad personality dimensions (Boyle, 2008). Second, FFM self-reports are sometimes subjective and may be influenced by many moderator factors such as cultures and situations (Boyle, 2008; “Five-Factor Model Personal. Across Cult.,” 2002). Some studies also show that many personality traits (such as openness to experience and conscientiousness) are not replicable across different samples (Hofstee, de Raad, & Goldberg, 1992; Johnson & Ostendorf, 1993). Despite all these limitations, the FFM has potentially been considered as a useful structure for describing the personality constructs. Moreover, in this paper, we have investigated the dynamic brain networks during resting-state. We believe that the use of cognitive tasks that stimulate the related networks for each personality trait may advance our understanding of individual differences in dynamic network features.

### Methodological considerations

First, in MEG analysis, the head model was computed from the individual MRI of each subject. Nevertheless, in EEG analysis, we used a template generated from MRIs of healthy controls, instead of a native MRI for EEG source connectivity. Recently, a study showed that there is no potential bias in the use of a template MRI as compared to individual MRI co-registration (Douw, Nieboer, Stam, Tewarie, & Hillebrand, 2018). In this context, a considerable number of EEG/MEG connectivity studies have used the template-based method due to the unavailability of native MRIs (Hassan et al., 2017; Kabbara et al., 2018; Lopez et al., 2014). However, we are aware that the use of subject-specific MRI is more recommended in clinical studies.

Second, we have adopted in each dataset the same pipeline (from data processing to networks construction) used by the previous studies dealing with the same datasets. Thus, for the EEG dataset, we used the wMNE/PLV combination to reconstruct the dynamic networks, as it is supported by two comparative studies (Mahmoud Hassan, Dufor, Merlet, Berrou, & Wendling, 2014; Mahmoud Hassan et al., 2016). For the MEG dataset, beamforming construction combined with amplitude correlation between band-limited power envelops was sustained by multiple studies (Brookes et al. 2012, Colclough et al. 2015, 2016; O’Neill et al. 2016).

Third, choosing the suitable window width is a crucial issue in constructing the dynamic functional networks. On the one hand, short windows do not contain sufficient information to accurately estimate connectivity. On the other hand, large windows may fail to capture the temporal changes of the brain networks. Hence, the ideal is to choose the shortest window that guarantees a sufficient number of data points over which the connectivity is calculated. This depends on the frequency band of interest that affects the degree of freedom in time series. In this study, we adapted the recommendation of Lachaux et al. (Lachaux et al., 2000) in selecting the smallest appropriate window length that is equal to where 6 is the number of ‘cycles’ at the given frequency band. The reproducibility of resting state results whilst changing the size of the sliding window was validated in a previous study (Kabbara et al., 2017a).

## Acknowledgments

This work was financed by the Rennes University, the Institute of Clinical Neuroscience of Rennes (Project named EEGCog) and AMU. The study was also funded by the National Council for Scientific Research (CNRS) in Lebanon. Authors would also like to thank the Lebanese Association for Scientific Research (LASER) for its support.

## Author contributions

Author contributions included conception and study design (VP, MH), data collection or acquisition (AW, VP), statistical analysis (AK, MH, JM), interpretation of results (AK, VP, MH, JM), drafting the manuscript work or revising it critically for important intellectual content (AK, MH, JM, VP) and approval of final version to be published and agreement to be accountable for the integrity and accuracy of all aspects of the work (All authors).

## References

Adelstein, J. S., Shehzad, Z., Mennes, M., DeYoung, C. G., Zuo, X. N., Kelly, C., … Milham, M. P. (2011). Personality is reflected in the brain’s intrinsic functional architecture. PLoS ONE. https://doi.org/10.1371/journal.pone.0027633

Aghajani, M., Veer, I. M., Van Tol, M. J., Aleman, A., Van Buchem, M. A., Veltman, D. J., … Van Der Wee, N. J. (2013). El neuroticismo y la extraversión están asociados con la conectividad funcional en estado de reposo de la amígdala. Cognitive, Affective and Behavioral Neuroscience. https://doi.org/10.3758/s13415-013-0224-0

Allen, E. A., Damaraju, E., Plis, S. M., Erhardt, E. B., Eichele, T., & Calhoun, V. D. (2014). Tracking whole-brain connectivity dynamics in the resting state. Cerebral Cortex, 24, 663–676. https://doi.org/10.1093/cercor/bhs352

Back, M. D., Schmukle, S. C., & Egloff, B. (2009). Predicting Actual Behavior From the Explicit and Implicit Self-Concept of Personality. Journal of Personality and Social Psychology. https://doi.org/10.1037/a0016229

Baker, A. P., Brookes, M. J., Rezek, I. A., Smith, S. M., Behrens, T., Smith, P. J. P., & Woolrich, M. (2014). Fast transient networks in spontaneous human brain activity. ELife, 2014. https://doi.org/10.7554/eLife.01867

Barrat, A., Barthélemy, M., Pastor-Satorras, R., & Vespignani, A. (2004). The architecture of complex weighted networks. Proceedings of the National Academy of Sciences of the United States of America, 101(11), 3747–3752. https://doi.org/10.1073/pnas.0400087101

Bassett, D. S., Porter, M. A., Wymbs, N. F., Grafton, S. T., Carlson, J. M., & Mucha, P. J. (2013). Robust detection of dynamic community structure in networks. Chaos, 23(1). https://doi.org/10.1063/1.4790830

Beaty, R. E., Kaufman, S. B., Benedek, M., Jung, R. E., Kenett, Y. N., Jauk, E., … Silvia, P. J. (2016a). Personality and complex brain networks: The role of openness to experience in default network efficiency. Human Brain Mapping, 37(2), 773–779. https://doi.org/10.1002/hbm.23065

Beaty, R. E., Kaufman, S. B., Benedek, M., Jung, R. E., Kenett, Y. N., Jauk, E., … Silvia, P. J. (2016b). Personality and complex brain networks: The role of openness to experience in default network efficiency. Human Brain Mapping. https://doi.org/10.1002/hbm.23065

Bey, K., Montag, C., Reuter, M., Weber, B., & Markett, S. (2015). Susceptibility to everyday cognitive failure is reflected in functional network interactions in the resting brain. NeuroImage. https://doi.org/10.1016/j.neuroimage.2015.07.041

Bland, j. M., & Altman, D. G. (1995). Multiple significance tests: The Bonferroni method. BMJ. https://doi.org/10.1136/bmj.310.6973.170

Blondel, V. D., Guillaume, J.-L., Lambiotte, R., & Lefebvre, E. (2008). Fast unfolding of communities in large networks. Journal of Statistical Mechanics: Theory and Experiment, 10008(10), 6. https://doi.org/10.1088/1742-5468/2008/10/P10008

Boyle, G. J. (2008). Critique of the five-factor model of personality. In The SAGE Handbook of Personality Theory and Assessment: Volume 1 - Personality Theories and Models. https://doi.org/10.4135/9781849200462.n14

Bressler, S. L. (1995). Large-scale cortical networks and cognition. Brain Research Reviews, Vol. 20, pp. 288–304. https://doi.org/10.1016/0165-0173(94)00016-I

Brookes, M. J., Woolrich, M. W., & Barnes, G. R. (2012). Measuring functional connectivity in MEG: A multivariate approach insensitive to linear source leakage. NeuroImage, 63(2), 910–920. https://doi.org/10.1016/j.neuroimage.2012.03.048

Brookes, Matthew J., Hale, J. R., Zumer, J. M., Stevenson, C. M., Francis, S. T., Barnes, G. R., … Nagarajan, S. S. (2011). Measuring functional connectivity using MEG: Methodology and comparison with fcMRI. NeuroImage, 56, 1082–1104. https://doi.org/10.1016/j.neuroimage.2011.02.054

Canli, T., & Amin, Z. (2002). Neuroimaging of emotion and personality: scientific evidence and ethical considerations. Brain and Cognition.

Colclough, G. L., Brookes, M. J., Smith, S. M., & Woolrich, M. W. (2015). A symmetric multivariate leakage correction for MEG connectomes. NeuroImage, 117, 439–448. https://doi.org/10.1016/j.neuroimage.2015.03.071

Colclough, G. L., Woolrich, M. W., Tewarie, P. K., Brookes, M. J., Quinn, A. J., & Smith, S. M. (2016). How reliable are MEG resting-state connectivity metrics? NeuroImage, 138, 284–293. https://doi.org/10.1016/j.neuroimage.2016.05.070

Cooper, N., Tompson, S., O’Donnell, M. B., & Falk, E. B. (2015). Brain activity in self- and value-related regions in response to online antismoking messages predicts behavior change. Journal of Media Psychology. https://doi.org/10.1027/1864-1105/a000146

Coquelet, N., De Tiège, X., Destoky, F., Roshchupkina, L., Bourguignon, M., Goldman, S., … Wens, V. (2020). Comparing MEG and high-density EEG for intrinsic functional connectivity mapping. NeuroImage, 116556. https://doi.org/10.1016/J.NEUROIMAGE.2020.116556

Costa, P. T., & McCrae, R. R. (1992). Neo PI-R professional manual. Psychological Assessment Resources. https://doi.org/10.1037/0003-066X.52.5.509

de Pasquale, F., Corbetta, M., Betti, V., & Della Penna, S. (2018). Cortical cores in network dynamics. NeuroImage. https://doi.org/10.1016/j.neuroimage.2017.09.063

de Pasquale, F., Penna, S. Della, Sporns, O., Romani, G. L., & Corbetta, M. (2015). A Dynamic Core Network and Global Efficiency in the Resting Human Brain. Cerebral Cortex, bhv185. https://doi.org/10.1093/cercor/bhv185

de Pasquale, Francesco, Della Penna, S., Snyder, A. Z., Marzetti, L., Pizzella, V., Romani, G. L., & Corbetta, M. (2012). A Cortical Core for Dynamic Integration of Functional Networks in the Resting Human Brain. Neuron, 74, 753–764. https://doi.org/10.1016/j.neuron.2012.03.031

Desikan, R. S., Sugonne, F., Fischl, B., Quinn, B. T., Dickerson, B. C., Blacker, D., … Killiany, R. J. (2006). An automated labeling system for subdividing the human cerebral cortex on MRI scans into gyral based regions of interest. NeuroImage, 31, 968–980. https://doi.org/10.1016/j.neuroimage.2006.01.021

Deyoung, C. G. (2006). Higher-order factors of the big five in a multi-informant sample. Journal of Personality and Social Psychology. https://doi.org/10.1037/0022-3514.91.6.1138

DeYoung, C. G. (2010). Personality Neuroscience and the Biology of Traits. Social and Personality Psychology Compass. https://doi.org/10.1111/j.1751-9004.2010.00327.x

Douw, L., Nieboer, D., Stam, C. J., Tewarie, P., & Hillebrand, A. (2018). Consistency of magnetoencephalographic functional connectivity and network reconstruction using a template versus native MRI for co-registration. Human Brain Mapping, 39(1), 104–119. https://doi.org/10.1002/hbm.23827

Dubois, J., Galdi, P., Han, Y., Paul, L. K., & Adolphs, R. (2018). Resting-State Functional Brain Connectivity Best Predicts the Personality Dimension of Openness to Experience. Personality Neuroscience. https://doi.org/10.1017/pen.2018.8

Edelman, G. M. (1993). Neural Darwinism: Selection and reentrant signaling in higher brain function. Neuron, Vol. 10, pp. 115–125. https://doi.org/10.1016/0896-6273(93)90304-A

Eickhoff, S. B., Stephan, K. E., Mohlberg, H., Grefkes, C., Fink, G. R., Amunts, K., & Zilles, K. (2005). A new SPM toolbox for combining probabilistic cytoarchitectonic maps and functional imaging data. NeuroImage, 25(4), 1325–1335. https://doi.org/10.1016/j.neuroimage.2004.12.034

Falk, E. B., O’Donnell, M. B., Tompson, S., Gonzalez, R., Dal Cin, S. D., Strecher, V., … An, L. (2015). Functional brain imaging predicts public health campaign success. Social Cognitive and Affective Neuroscience. https://doi.org/10.1093/scan/nsv108

Furr, R. M. (2009). Personality psychology as a truly behavioural science. European Journal of Personality. https://doi.org/10.1002/per.724

Fuster, J. M. (2010). Cortex and Mind: Unifying Cognition. In Cortex and Mind: Unifying Cognition. https://doi.org/10.1093/acprof:oso/9780195300840.001.0001

Gao, Q. (2013). Erratum: Extraversion and neuroticism relate to topological properties of resting-state brain networks. Frontiers in Human Neuroscience. https://doi.org/10.3389/fnhum.2013.00448

Girvan, M., & Newman, M. E. J. (2002). Community structure in social and biological networks. Proceedings of the National Academy of Sciences of the United States of America, 99(12), 7821–7826. https://doi.org/10.1073/pnas.122653799

Goldman-Rakic, P. S. (1988). Topography of Cognition: Parallel Distributed Networks in Primate Association Cortex. Annual Review of Neuroscience, 11(1), 137–156. https://doi.org/10.1146/annurev.ne.11.030188.001033

Gramfort, A., Papadopoulo, T., Olivi, E., & Clerc, M. (2010). OpenMEEG: opensource software for quasistatic bioelectromagnetics. Biomedical Engineering Online, 9, 45. https://doi.org/10.1186/1475-925X-9-45

Gray, J. C., Owens, M. M., Hyatt, C. S., & Miller, J. D. (2018). No evidence for morphometric associations of the amygdala and hippocampus with the five-factor model personality traits in relatively healthy young adults. PLoS ONE. https://doi.org/10.1371/journal.pone.0204011

Greicius, M. D., Krasnow, B., Reiss, A. L., & Menon, V. (2003). Functional connectivity in the resting brain: a network analysis of the default mode hypothesis. Proceedings of the National Academy of Sciences of the United States of America, 100, 253–258. https://doi.org/10.1073/pnas.0135058100

Hamalainen, M. S., & Ilmoniemi, R. J. (1994). Interpreting magnetic fields of the brain: minimum norm estimates. Medical & Biological Engineering & Computing, 32(1), 35–42. https://doi.org/10.1007/BF02512476

Hassan, M., Chaton, L., Benquet, P., Delval, A., Leroy, C., Plomhause, L., … Dujardin, K. (2017). Functional connectivity disruptions correlate with cognitive phenotypes in Parkinson’s disease. NeuroImage: Clinical, 14, 591–601. https://doi.org/10.1016/j.nicl.2017.03.002

Hassan, M, & Wendling, F. (2018). Electroencephalography source connectivity: toward high time / space resolution brain networks. IEEE Signal Processing Magazine, 1–25.

Hassan, Mahmoud, Dufor, O., Merlet, I., Berrou, C., & Wendling, F. (2014). EEG source connectivity analysis: From dense array recordings to brain networks. PLoS ONE, 9. https://doi.org/10.1371/journal.pone.0105041

Hassan, Mahmoud, Merlet, I., Mheich, A., Kabbara, A., Biraben, A., Nica, A., & Wendling, F. (2016). Identification of Interictal Epileptic Networks from Dense-EEG. Brain Topography, pp. 1–17. https://doi.org/10.1007/s10548-016-0517-z

Hong, R. Y., Paunonen, S. V., & Slade, H. P. (2008). Big Five personality factors and the prediction of behavior: A multitrait-multimethod approach. Personality and Individual Differences. https://doi.org/10.1016/j.paid.2008.03.015

Iacoboni, M., & Zaidel, E. (2004). Interhemispheric visuo-motor integration in humans: The role of the superior parietal cortex. Neuropsychologia. https://doi.org/10.1016/j.neuropsychologia.2003.10.007

Jaccard, J. J. (1974). Predicting social behavior from personality traits. Journal of Research in Personality. https://doi.org/10.1016/0092-6566(74)90057-9

Kabbara, A., Eid, H., El Falou, W., Khalil, M., Wendling, F., & Hassan, M. (2018). Reduced integration and improved segregation of functional brain networks in Alzheimer’s disease. Journal of Neural Engineering, 15(2). https://doi.org/10.1088/1741-2552/aaaa76

Kabbara, A., Eid, H., Falou, E. L., Khalil, M., Wendling, F., & Hassan, M. (2018). Reduced integration and improved segregation of functional brain networks in Alzheimer’s disease. Journal of Neural Engineeri.

Kabbara, A., Falou, W. E. L., Khalil, M., Wendling, F., & Hassan, M. (2017a). The dynamic functional core network of the human brain at rest. Scientific Reports, 7(1), 2936.

Kabbara, A., Falou, W. E. L., Khalil, M., Wendling, F., & Hassan, M. (2017b). The dynamic functional core network of the human brain at rest. (August 2016), 1–16. https://doi.org/10.1038/s41598-017-03420-6

Kabbara, A., Khalil, M., O’Neill, G., Dujardin, K., El Traboulsi, Y., Wendling, F., & Hassan, M. (2019). Detecting modular brain states in rest and task. Network Neuroscience. https://doi.org/10.1162/netn_a_00090

Kenett, Y. N., Betzel, R. F., & Beaty, R. E. (2020). Community structure of the creative brain at rest. NeuroImage, 210, 116578. https://doi.org/10.1016/J.NEUROIMAGE.2020.116578

Knyazev, G. G. (2012). EEG delta oscillations as a correlate of basic homeostatic and motivational processes. Neuroscience and Biobehavioral Reviews. https://doi.org/10.1016/j.neubiorev.2011.10.002

Kosslyn, S. (2007). Cognitive Psychology: Mind and Brain. New Jersey: Prentice Hall.

Kyeong, S., Kim, E., Park, H. J., & Hwang, D. U. (2014). Functional network organizations of two contrasting temperament groups in dimensions of novelty seeking and harm avoidance. Brain Research. https://doi.org/10.1016/j.brainres.2014.05.037

Lachaux, J.-P., Rodriguez, E., Le van Quyen, M., Lutz, A., Martinerie, J., & Varela, F. J. (2000). Studying single-trials of phase synchronous activity in the brain. International Journal of Bifurcation and Chaos, 10(10), 2429–2439. https://doi.org/10.1142/S0218127400001560

Larson-Prior, L. J., Oostenveld, R., Della Penna, S., Michalareas, G., Prior, F., Babajani-Feremi, A., … Snyder, A. Z. (2013). Adding dynamics to the Human Connectome Project with MEG. NeuroImage. https://doi.org/10.1016/j.neuroimage.2013.05.056

Li, T., Yan, X., Li, Y., Wang, J., Li, Q., Li, H., & Li, J. (2017). Neuronal correlates of individual differences in the big five personality traits: Evidences from cortical morphology and functional homogeneity. Frontiers in Neuroscience, 11 (JUL), 1–8. https://doi.org/10.3389/fnins.2017.00414

Liu, W. Y., Weber, B., Reuter, M., Markett, S., Chu, W. C., & Montag, C. (2013). The Big Five of Personality and structural imaging revisited: A VBM - DARTEL study. NeuroReport. https://doi.org/10.1097/WNR.0b013e328360dad7

Lopez, M. E., Bruna, R., Aurtenetxe, S., Pineda-Pardo, J. A., Marcos, A., Arrazola, J., … Maestu, F. (2014). Alpha-Band Hypersynchronization in Progressive Mild Cognitive Impairment: A Magnetoencephalography Study. Journal of Neuroscience, 34(44), 14551–14559. https://doi.org/10.1523/JNEUROSCI.0964-14.2014

Markett, S., Weber, B., Voigt, G., Montag, C., Felten, A., Elger, C., & Reuter, M. (2013). Intrinsic connectivity networks and personality: The temperament dimension harm avoidance moderates functional connectivity in the resting brain. Neuroscience. https://doi.org/10.1016/j.neuroscience.2013.02.056

Markett, Sebastian, Montag, C., Melchers, M., Weber, B., & Reuter, M. (2016). Anxious personality and functional efficiency of the insular-opercular network: A graph-analytic approach to resting-state fMRI. Cognitive, Affective and Behavioral Neuroscience. https://doi.org/10.3758/s13415-016-0451-2

Markett, Sebastian, Montag, C., & Reuter, M. (2018). Network Neuroscience and Personality. Personality Neuroscience. https://doi.org/10.1017/pen.2018.12

McAdams, D. P. (1992). The Five-Factor Model In Personality: A Critical Appraisal. Journal of Personality. https://doi.org/10.1111/j.1467-6494.1992.tb00976.x

McCrae, R. R., & John, O. P. (1992). An Introduction to the Five-Factor Model and Its Applications. Journal of Personality. https://doi.org/10.1111/j.1467-6494.1992.tb00970.x

Mesulam, M. -M. (1990). Large-scale neurocognitive networks and distributed processing for attention, language, and memory. Annals of Neurology, 28(5), 597–613. https://doi.org/10.1002/ana.410280502

Mulders, P., Llera, A., Tendolkar, I., van Eijndhoven, P., & Beckmann, C. (2018). Personality Profiles Are Associated with Functional Brain Networks Related to Cognition and Emotion. Scientific Reports, 8(1), 1–8. https://doi.org/10.1038/s41598-018-32248-x

O’Neill, G. C., Tewarie, P. K., Colclough, G. L., Gascoyne, L. E., Hunt, B. A. E., Morris, P. G., … Brookes, M. J. (2016). Measurement of Dynamic Task Related Functional Networks using MEG. NeuroImage, in press. https://doi.org/10.1016/j.neuroimage.2016.08.061

O’Neill, G. C., Tewarie, P., Vidaurre, D., Liuzzi, L., Woolrich, M. W., & Brookes, M. J. (2017). Dynamics of large-scale electrophysiological networks: A technical review. NeuroImage. https://doi.org/10.1016/j.neuroimage.2017.10.003

Omura, K., Constable, R. T., & Canli, T. (2005). Amygdala gray matter concentration is associated with extraversion and neuroticism. NeuroReport. https://doi.org/10.1097/01.wnr.0000186596.64458.76

Ormel, J., Bastiaansen, A., Riese, H., Bos, E. H., Servaas, M., Ellenbogen, M., … Aleman, A. (2013). The biological and psychological basis of neuroticism: Current status and future directions. Neuroscience and Biobehavioral Reviews. https://doi.org/10.1016/j.neubiorev.2012.09.004

Paban, V., Deshayes, C., Ferrer, M.-H., Weill, A., & Alescio-Lautier, B. (2018). Resting Brain Functional Networks and Trait Coping. Brain Connectivity. https://doi.org/10.1089/brain.2018.0613

Pelphrey, K. A., & Carter, E. J. (2008). Brain mechanisms for social perception: Lessons from autism and typical development. Annals of the New York Academy of Sciences. https://doi.org/10.1196/annals.1416.007

Pelphrey, K. A., & Morris, J. P. (2006). Brain mechanisms for interpreting the actions of others from biological-motion cues. Current Directions in Psychological Science. https://doi.org/10.1111/j.0963-7214.2006.00423.x

Plaisant, O., Courtois, R., Réveillère, C., Mendelsohn, G. A., & John, O. P. (2010). Validation par analyse factorielle du Big Five Inventory français (BFI-Fr). Analyse convergente avec le NEO-PI-R. Annales Medico-Psychologiques. https://doi.org/10.1016/j.amp.2009.09.003

Riccelli, R., Toschi, N., Nigro, S., Terracciano, A., & Passamonti, L. (2017). Surface-based morphometry reveals the neuroanatomical basis of the five-factor model of personality. Social Cognitive and Affective Neuroscience. https://doi.org/10.1093/scan/nsw175

Rizkallah, J., Benquet, P., Kabbara, A., Dufor, O., Wendling, F., & Hassan, M. (2018). Dynamic reshaping of functional brain networks during visual object recognition. Journal of Neural Engineering. https://doi.org/10.1088/1741-2552/aad7b1

Rubinov, M., & Sporns, O. (2011). Weight-conserving characterization of complex functional brain networks. NeuroImage, 56(4), 2068–2079. https://doi.org/10.1016/j.neuroimage.2011.03.069

Sawada, R., Sato, W., Uono, S., Kochiyama, T., Kubota, Y., Yoshimura, S., & Toichi, M. (2016). Neuroticism delays detection of facial expressions. PLoS ONE. https://doi.org/10.1371/journal.pone.0153400

Singer, T. (2006). The neuronal basis and ontogeny of empathy and mind reading: Review of literature and implications for future research. Neuroscience and Biobehavioral Reviews. https://doi.org/10.1016/j.neubiorev.2006.06.011

Sporns, O, Chialvo, D. R., Kaiser, M., & Hilgetag, C. C. (2004). Organization, development and function of complex brain networks. Trends In Cognitive Sciences, 8(9), 418–425. https://doi.org/10.1016/j.tics.2004.07.008

Sporns, Olaf, & Betzel, R. F. (2016). Modular Brain Networks. Annual Review of Psychology, 67(1), 613–640. https://doi.org/10.1146/annurev-psych-122414-033634

Suslow, T., Kugel, H., Reber, H., Bauer, J., Dannlowski, U., Kersting, A., … Egloff, B. (2010). Automatic brain response to facial emotion as a function of implicitly and explicitly measured extraversion. Neuroscience. https://doi.org/10.1016/j.neuroscience.2010.01.038

Tadel, F., Baillet, S., Mosher, J. C., Pantazis, D., & Leahy, R. M. (2011). Brainstorm: A user-friendly application for MEG/EEG analysis. Computational Intelligence and Neuroscience, 2011. https://doi.org/10.1155/2011/879716

Terracciano, A. (2003). The Italian version of the NEO PI-R: Conceptual and empirical support for the use of targeted rotation. Personality and Individual Differences. https://doi.org/10.1016/S0191-8869(03)00035-7

The Five-Factor Model of Personality Across Cultures. (2002). In The Five-Factor Model of Personality Across Cultures. https://doi.org/10.1007/978-1-4615-0763-5

Tian, F., Wang, J., Xu, C., Li, H., & Ma, X. (2018). Focusing on the differences of resting-state brain networks, using a data-driven approach to explore the functional neuroimaging characteristics of extraversion trait. Frontiers in Neuroscience, 12(MAR), 1–8. https://doi.org/10.3389/fnins.2018.00109

Tomeček, D., & Androvičová, R. (2017). Personality Reflection in the Brain’s Intrinsic Functional Architecture Remains Elusive. Neuroimage.

Tompson, S. H., Falk, E. B., Vettel, J. M., & Bassett, D. S. (2018). Network Approaches to Understand Individual Differences in Brain Connectivity: Opportunities for Personality Neuroscience. Personality Neuroscience. https://doi.org/10.1017/pen.2018.4

Toschi, N., Riccelli, R., Indovina, I., Terracciano, A., & Passamonti, L. (2018). Functional Connectome of the Five-Factor Model of Personality. Personality Neuroscience, 1. https://doi.org/10.1017/pen.2017.2

Traud, A. L., Kelsic, E. D., Mucha, P. J., & Porter, M. A. (2008). Comparing Community Structure to Characteristics in Online Collegiate Social Networks. 53(3), 526–543. https://doi.org/10.1137/080734315

Van Essen, D. C., Ugurbil, K., Auerbach, E., Barch, D., Behrens, T. E. J., Bucholz, R., … Yacoub, E. (2012). The Human Connectome Project: A data acquisition perspective. NeuroImage. https://doi.org/10.1016/j.neuroimage.2012.02.018

Van Veen, B. D., Van Drongelen, W., Yuchtman, M., & Suzuki, A. (1997). Localization of brain electrical activity via linearly constrained minimum variance spatial filtering. IEEE Transactions on Biomedical Engineering. https://doi.org/10.1109/10.623056

Yarkoni, T. (2014). Neurobiological substrates of personality: A critical overview. In APA handbook of personality and social psychology, Volume 4: Personality processes and individual differences. https://doi.org/10.1037/14343-003

Zelenski, J. M., & Larsen, R. J. (1999). Susceptibility to affect: A comparison of three personality taxonomies. Journal of Personality. https://doi.org/10.1111/1467-6494.00072

